# Transcriptomic analysis of ribosome biogenesis and pre-rRNA processing during growth stress in *Entamoeba histolytica*

**DOI:** 10.1101/2021.08.01.454488

**Authors:** Sarah Naiyer, Shashi Shekhar Singh, Devinder Kaur, Yatendra Pratap Singh, Amartya Mukherjee, Alok Bhattacharya, Sudha Bhattacharya

## Abstract

Ribosome biogenesis, a multi-step process involving the transcription, modification, folding and processing of rRNA is the major consumer of cellular energy. It involves the sequential assembly of ribosomal proteins (RP)s via more than 200 ribogenesis factors. Unlike model organisms where transcription of rRNA and RP genes slows down during stress, in *Entamoeba histolytica*, pre-rRNA synthesis continues, and unprocessed pre-rRNA accumulates. To gain insight into the vast repertoire of ribosome biogenesis factors and understand the major components playing role during stress we computationally identified the ribosome biogenesis factors in *E. histolytica.* Of the total ∼279 *S. cerevisiae* proteins, we could only find 188 proteins in *E. histolytica*. Some of the proteins missing in *E. histolytica* were also missing in humans. A number of proteins represented by multiple genes in *S. cerevisiae* had only a single copy in *E. histolytica.* It was interesting to note that *E. histolytica* lacked mitochondrial ribosome biogenesis factors and had far less RNase components as compared to *S. cerevisiae*. Northern hybridization using probes from different spacer regions depicted the accumulation of unprocessed intermediates during stress. Transcriptomic studies revealed the differential regulation of a number of ribosomal factors both in serum-starved and RRP6KD conditions. The ARB1 protein involved at multiple steps of ribosome biogenesis and NEP1 and TSR3 involved in chemical modification of 18S rRNA previously shown to accumulate pre-rRNA precursors upon downregulation in *S. cerevisiae* and humans were included. The data reveals the importance of some of the major factors required for regulating pre-rRNA processing during stress. This is the first report on the complete repertoire of ribosome biogenesis factors in *E. histolytica*.

## Introduction

The ribosome, in addition to being a conserved molecular factory for protein synthesis, is a complex multifaceted machinery engaged in the spatiotemporal control of gene expression (1). Due to the very large number of ribosomes in a typical cell their biosynthesis consumes a major part of cellular energy and nuclear space (2,3). Transcription of ribosomal RNA genes, and ribosome formation is thus highly regulated in response to general metabolism and specific environmental conditions (4,5). In eukaryotes, the small subunit of ribosome (40S) is composed of 18S rRNA assembled with 33 ribosomal proteins (RPs), while the large subunit (60S) has 5S, 5.8S, and 25S/28S rRNAs associated with 47 RPs (6). Ribosome biogenesis takes place in the nucleolus, in a multi-step process involving the transcription, modification, folding and processing of rRNA along with the assembly of RPs with the help of more than 200 ribogenesis factors. Much has been learnt about this complex process in model systems, especially *Saccharomyces cerevisiae*, and has been reviewed in various authoritative articles (7–13). A brief description of the ribosomal biogenesis process is as follows.

The primary rRNA transcript (35S in *S. cerevisiae*) undergoes multiple specific cleavages at the 5’ and 3’ external transcribed spacers (5’-ETS and 3’-ETS) and the internal transcribed spacers 1 (ITS1) and 2 (ITS2), to generate the mature 18S, 5.8S, and 25S/28S rRNAs (14). Pre-rRNA processing begins co-transcriptionally by the formation of 90S SSU processome, a ribonucleoprotein complex of ∼70 assembly factors and several small nucleolar RNAs (snoRNAs). The protein components of the processome include RNA-binding proteins, endonucleases, RNA helicases, ATPases, GTPases, and kinases. The SSU processome proteins identified by co-purification with U3 snoRNA are termed the U three proteins (UTPs) (15). Both the RNA and protein components of the SSU processome are considered to function as a pre-rRNA chaperone to assist in the correct folding of nascent pre-rRNA. This enables sequence-specific cleavages at sites A0, A1 and A2 to generate the pre-18S rRNA. Amongst the snoRNAs, U3 is a non-canonical C/D box-containing snoRNA that has the specialized function of assisting in pre-rRNA processing and maturation (16). It associates with the canonical C/D box-specific proteins, and the U3 snoRNA-specific RRP9/U3-55K protein (17) to form U3snoRNPs that base pair at specific sites within the 5’-ETS and the pre-18S rRNA to carry out processing at sites A0, A1 and A2 (18). The rRNA precursor is also chemically modified by the canonical C/D-box and the H/ACA-box containing snoRNAs required, respectively, for methylation and pseudo-uridylation of the rRNAs (19–22). After the modification, the snoRNAs that are base paired with their target sites in the pre-rRNA, are removed by the action of DExH/H-box RNA helicases. The correctly folded pre-rRNAs are assembled with RPs, which along with biogenesis factors are synthesized in the cytoplasm, and transported to the nucle(ol)us for assembly of pre-ribosomal particles (23).

The initial cleavages at sites A0 and A1 in the 5’-ETS lead to the 90S pre-ribosomal particle being separated into pre-40S and pre-60S particles via cleavage at site A2. It requires the release of 5’-ETS rRNA that is complexed with UTP-A, UTP-B, and U3snoRNPs (24). The endoribonucleases encoded by UTP24 (25), and RCL1 are likely involved in cleavage at A2 (26). The RNA helicase DHR1 also plays an important role in the dismantling of the 90S particle. After cleavage and trimming, the pre-40S particle is translocated to the cytoplasm through the nuclear pore complex. This process is facilitated by the GTPases GSP1/RAN and CRM1/XPO1 (27). The final stage of pre-40S maturation in the cytoplasm includes the NOB1 endoribonuclease-catalyzed cleavage at site D of the 20S pre-rRNA to form the mature 18S-rRNA (28).

Compared with 40S, the assembly of 60S particle appears to be much more complex. 5.8S rRNAs with different 5’-ends are produced by two alternative pathways (29,30). The maturation of 5’-end of 5.8S rRNA is coordinated with cleavage of the 3’-end of 25S/28S rRNA, which is a prerequisite for endonucleolytic cleavage of ITS2 at site C2 (31). Following this cleavage, the 7S precursor is released. The maturation of 5.8S 3’-end from the 7S precursor involves the nuclear exosome, assisted by the RNA helicase DOB1/MTR4, and 3’-5’ exonucleases including RRP6 (32,33). The exonuclease RAT1 generates the mature 5’-end of the 25S rRNA from 26S precursor (34). Processing at both ends of ITS2 is catalyzed by the protein complex LAS1 (35). A number of export factors are required for final translocation of pre-60S particles to the cytoplasm. These factors interact with specific sites of the pre-60S ribosome. The export adaptor NMD3 binds to the large subunit and recruits the exportin CRM1/ XPO1 via its NES sequence (36). Other export factors include the MEX67-MTR2 heterodimer, and factors that shield the charged ribosomal surface against hydrophobic environment within the NPC channel (37,38).

Although many of the ribosomal biogenesis components described above are well conserved in yeast and human, a number of yeast-specific proteins have no known human homologs, pointing to important functional diversity amongst organisms. It will be interesting to explore the variations in this highly conserved function in evolutionarily distant organisms. We have been investigating the regulation of pre-rRNA processing and ribosome biogenesis under growth stress in the primitive parasitic protist, *Entamoeba histolytica*, which causes amoebiasis in humans (39). In this organism the rRNA genes are located exclusively on extrachromosomal circular molecules (40,41), and the nucleolus is organized at the nuclear periphery (42). Each rDNA circle may contain either one copy of the rDNA transcription unit, or two copies organized as inverted repeats. In *E. histolytica* strain HM-1:IMSS, the 14 kb rDNA circle designated EhR2 contains one rDNA unit (43), for which the transcription start point has been mapped (44–46). We have earlier shown that pre-rRNA processing was inhibited in *E. histolytica* cells subjected to growth stress by serum starvation, leading to accumulation of unprocessed pre-rRNAs. There was strong accumulation of partially processed fragments of the 5’-ETS, which are otherwise rapidly degraded in unstressed cells (46). Anomalies in pre-rRNA processing during growth stress could result from downregulation of various components of the processing machinery, including endoribonucleases, exoribonucleases, and helicases. Our previous work has shown that the exosome-associated 3’-5’ exoribonuclease EhRRP6 is down-regulated in serum-starved *E. histolytica*, and the enzyme is lost from the nuclei in starved cells (47). Since this enzyme is known to be involved in the removal of 5’-ETS sub fragments in model organisms (48,49), its down regulation during serum starvation could lead to the accumulation of partially processed fragments of 5’-ETS in *E. histolytica*.

To obtain a more comprehensive picture of pre-rRNA processing, we have now computationally identified the *S. cerevisiae* homologs of pre-rRNA processing and ribosome biogenesis factors in *E. histolytica.* Using northern hybridization, we have shown the accumulation of unprocessed intermediates during serum starvation. Further, we have used transcriptomic analysis to study the regulation of these components during growth stress, and in EhRRP6 down-regulated cell lines. The data provide insights into the unique features of ribosome biogenesis components in *E. histolytica*.

## Results and Discussion

### Computational identification of *E. histolytica* ribosome biogenesis and pre-rRNA processing proteins

Ribosome biogenesis factors in eukaryotes affect diverse cellular pathways, and perturbations in ribosome biogenesis are linked to human disease (50). To understand the status of ribosome biogenesis in *E. histolytica*, we searched the *E. histolytica* sequence database for presence of all matches with proteins known to be involved in ribosome biogenesis. Since *S. cerevisiae* is the model organism giving the deepest insights in ribosome biogenesis, we enlisted all *S. cerevisiae* proteins involved in the process and looked for their counterparts in *E. histolytica* (as detailed in methods section). A list of the classes of proteins known to play a role in ribosome biogenesis is shown in Table 1. The complete list of proteins is given in Supplementary Table 1. Of the total ∼279 *S. cerevisiae* proteins listed (51), we could find matches for 188 proteins in *E. histolytica* (Table 1). Their transcription, as determined from our RNA-Seq data (52,53) ranged from very high to low.

**Table 1:** Ribosome biogenesis factors and their expression status in E. histolytica.

| Classes of proteins involved in various steps of ribosome biogenesis |  | No of genes in Sc | No of genes in Eh | Expression level (VH,H>M>VL,L>A) |
| --- | --- | --- | --- | --- |
| **SnoRNPs** | Box C/D | 4 | 5 | 03>2>0>0 |
|  | Box H/ACA | 6 | 8 | 07>1>0>0 |
| **90s particles** | U3 snoRNA associated proteins | 30 | 27 | 0>24>2>1 |
|  | Helicase | 9 | 18 | 1>13>03>1 |
|  | RNase | 1 | 1 | 0>0>0>1 |
|  | GTPase | 3 | 1 | 0>1>0>0 |
|  | Kinase | 4 | 6 | 0>5>0>1 |
|  | Other 90S particle | 22 | 20 | 0>13>5>2 |
| **Pre-40S particles** | Kinases | 3 | 7 | 0>6>1>0 |
|  | Export and cytoplasmic maturation factors | 20 | 22 | 1>17>4>0 |
|  | Other pre-40S particles | 6 | 13 | 2>7>3>1 |
| **Pre-60S particles** | Helicase | 11 | 13 | 0>11>2>0 |
|  | RNase | 15 | 5 | 0>5>0>0 |
|  | GTPase | 5 | 7 | 3>4>0>0 |
|  | Kinase | 2 | -- |  |
|  | Export and cytoplasmic maturation factors | 22 | 23 | 3>11>4>5 |
|  | 5-FMC complex | 3 | 2 | 0>2>0>0 |
|  | Other pre-60S particle | 52 | 39 | 2>34>3>0 |
| **Other ribosome biogenesis factors** |  | 13 | 6 | 0>6>0>0 |
| **Miscellaneous proteins** |  | 54 | 41 | 15>22>0>4>0 |

\*VH, very high; H, high; M, medium; L, low; A, absent.

Of the 188 listed *E. histolytica* genes involved in pre-rRNA processing, transcripts of 47 genes were either absent or they belonged to the very low/ low-expression categories. The low-expressing genes mainly included helicases, some kinases, phospholipid transporting P-type ATPase, rRNA biogenesis protein RRP5, exosome complex exonuclease RRP44, tRNA splicing endonuclease and some HSP70 family genes. Around 44 genes were associated with high/ very high expression levels and included the U3snoRNA family, fibrillarin, 13kDa ribonucleoprotein-associated protein, centromere/microtubule binding protein, snoRNP protein GAR1, H/ACA ribonucleoprotein complex subunit 2-like protein. These protein sets are well established to play the pivotal roles in pre-rRNA processing. Other protein families included protein phosphatase, elongation factor 2, snRNP Sm D2, LSM domain containing protein, enhancer binding protein 2 (EBP2), eukaryotic translation initiation factor 6 and 40S ribosomal proteins S2, S4, S6, S7, S9, S21. We could not detect 91 proteins in *E. histolytica*, some of which play an essential role in *S. cerevisiae*. Some of these were also missing in humans, suggesting their divergent nature (Table 2). The evolution of alternative pathways in *S. cerevisiae* or functional redundancy in *E. histolytica* may account for this loss. Alternatively, some of these proteins might have been missed due to poor sequence homology.

**Table 2:**
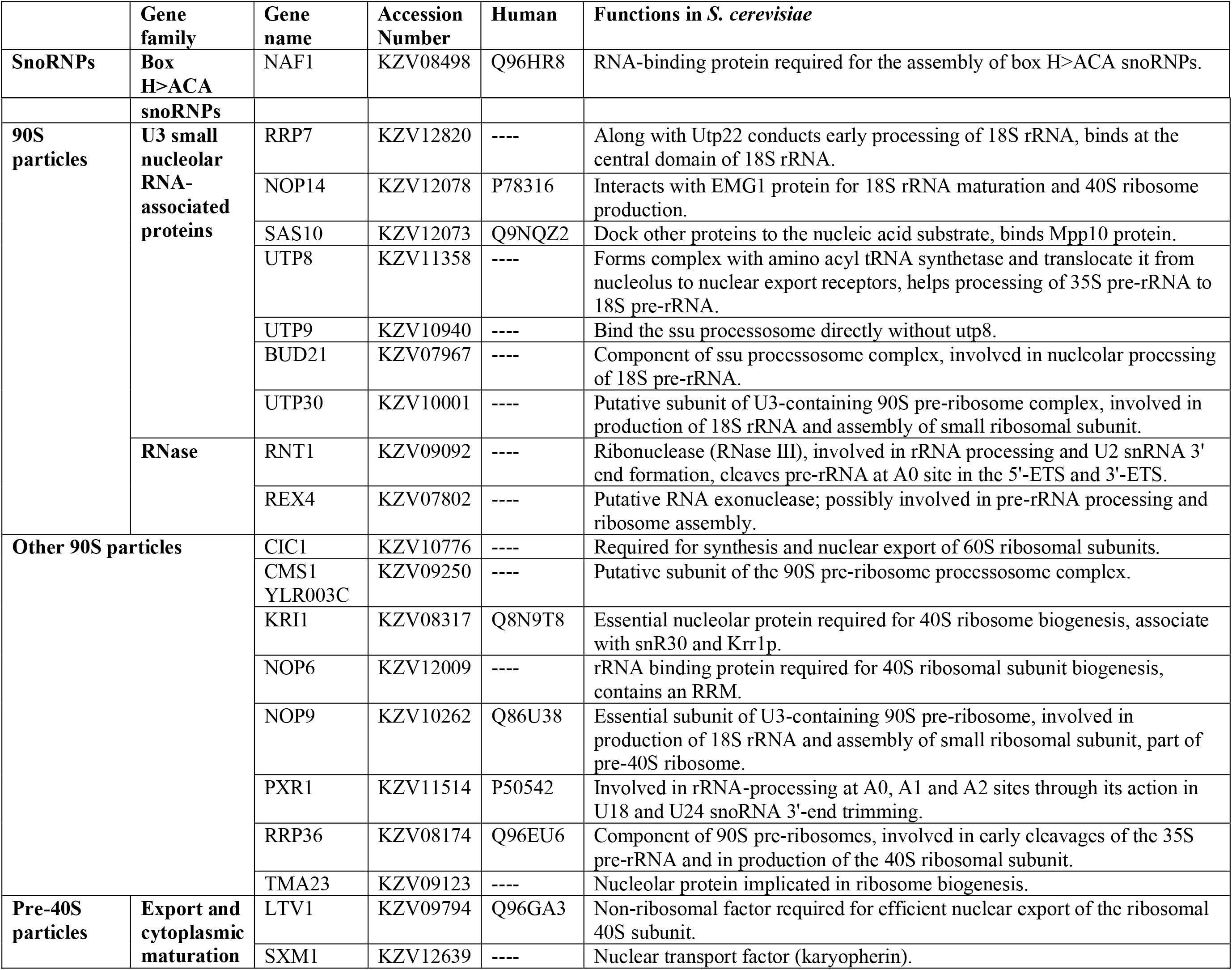
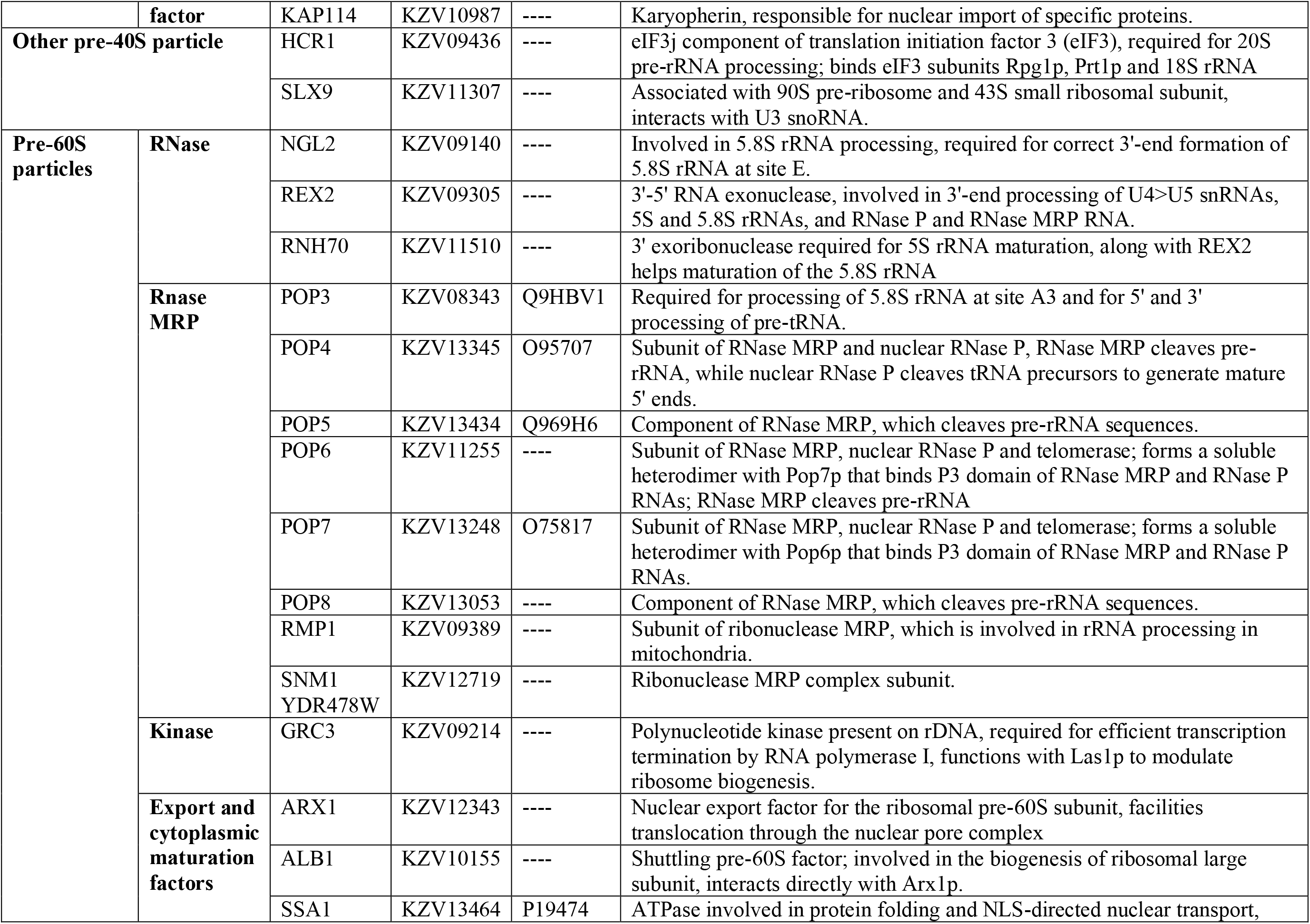
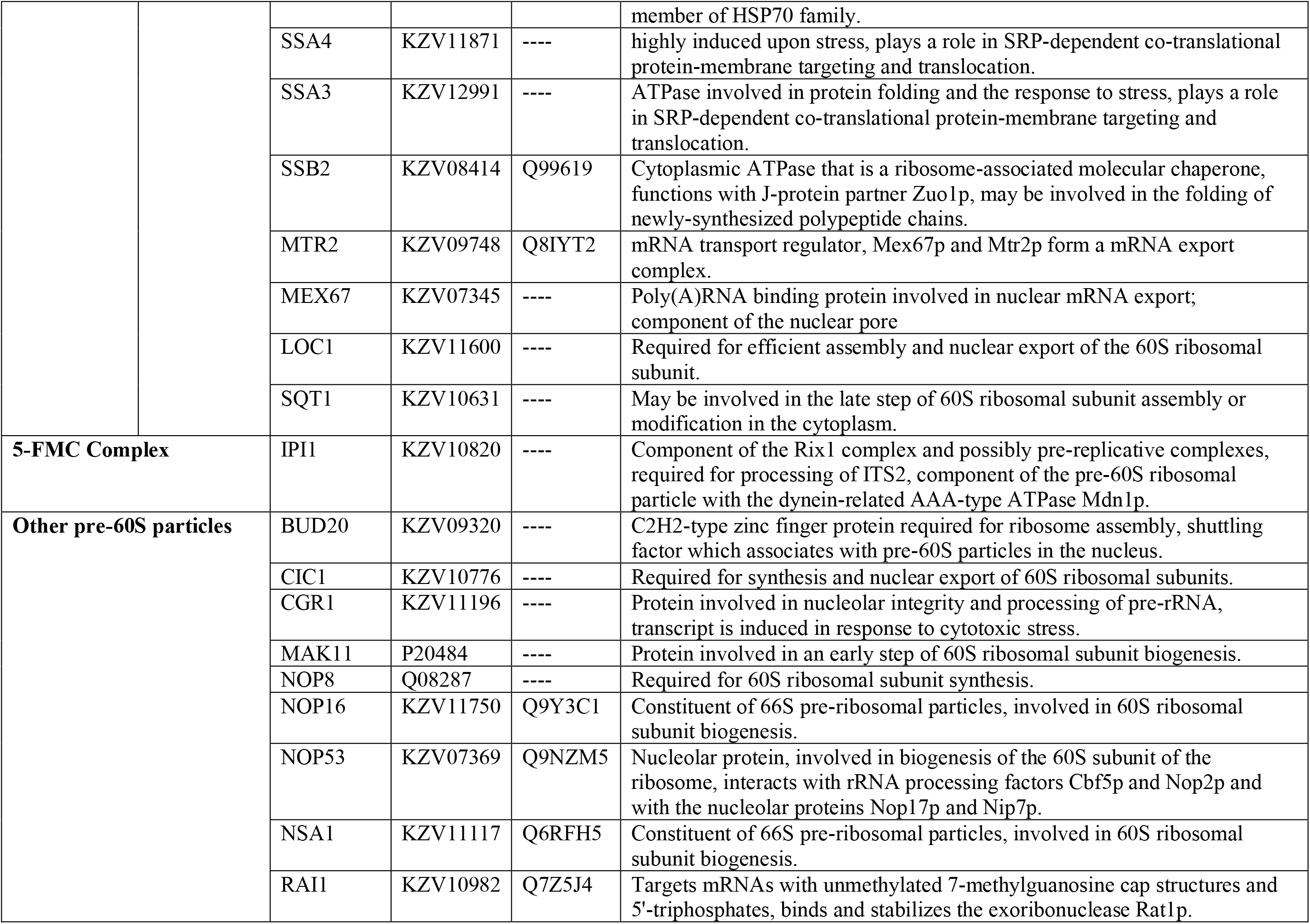
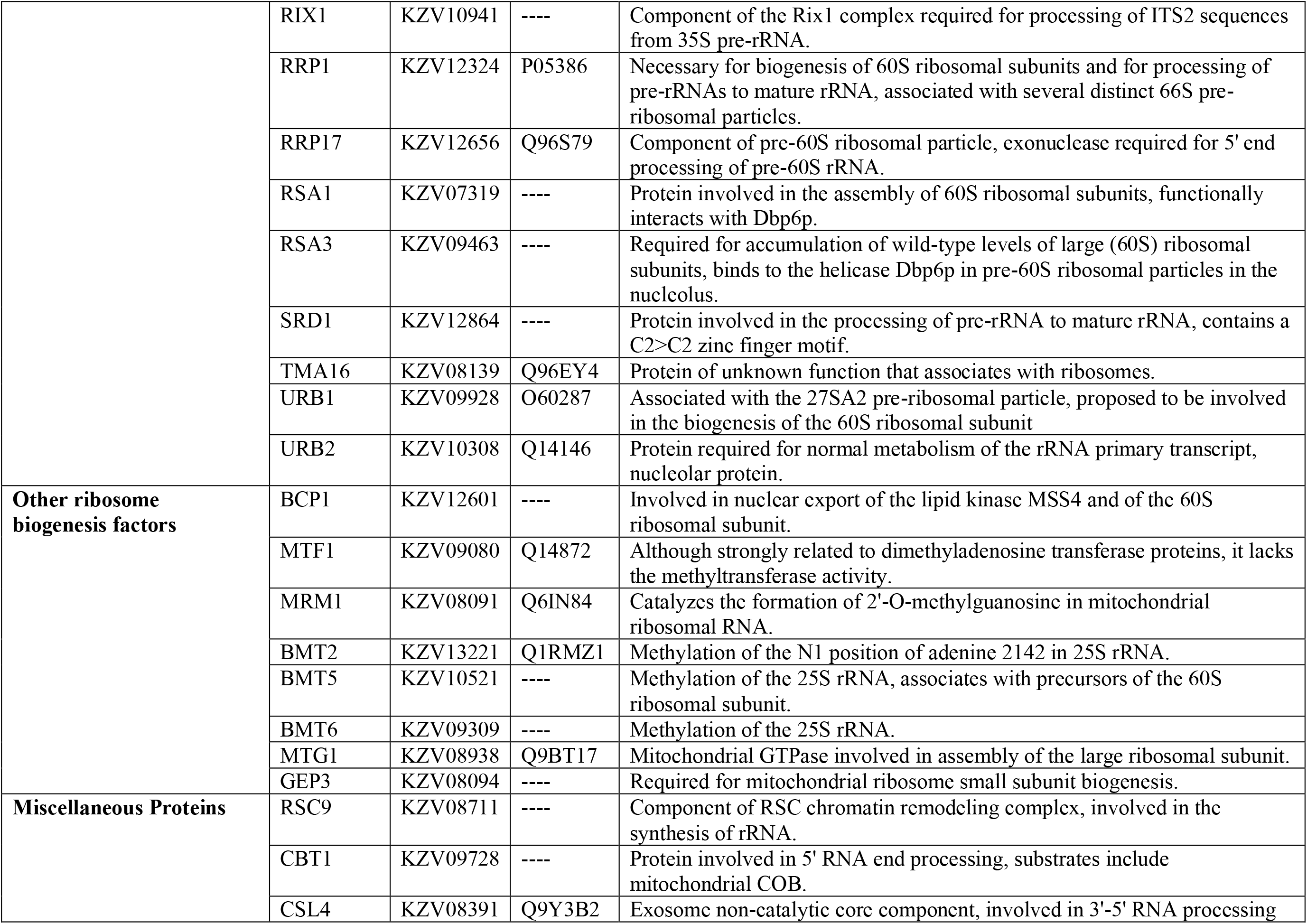
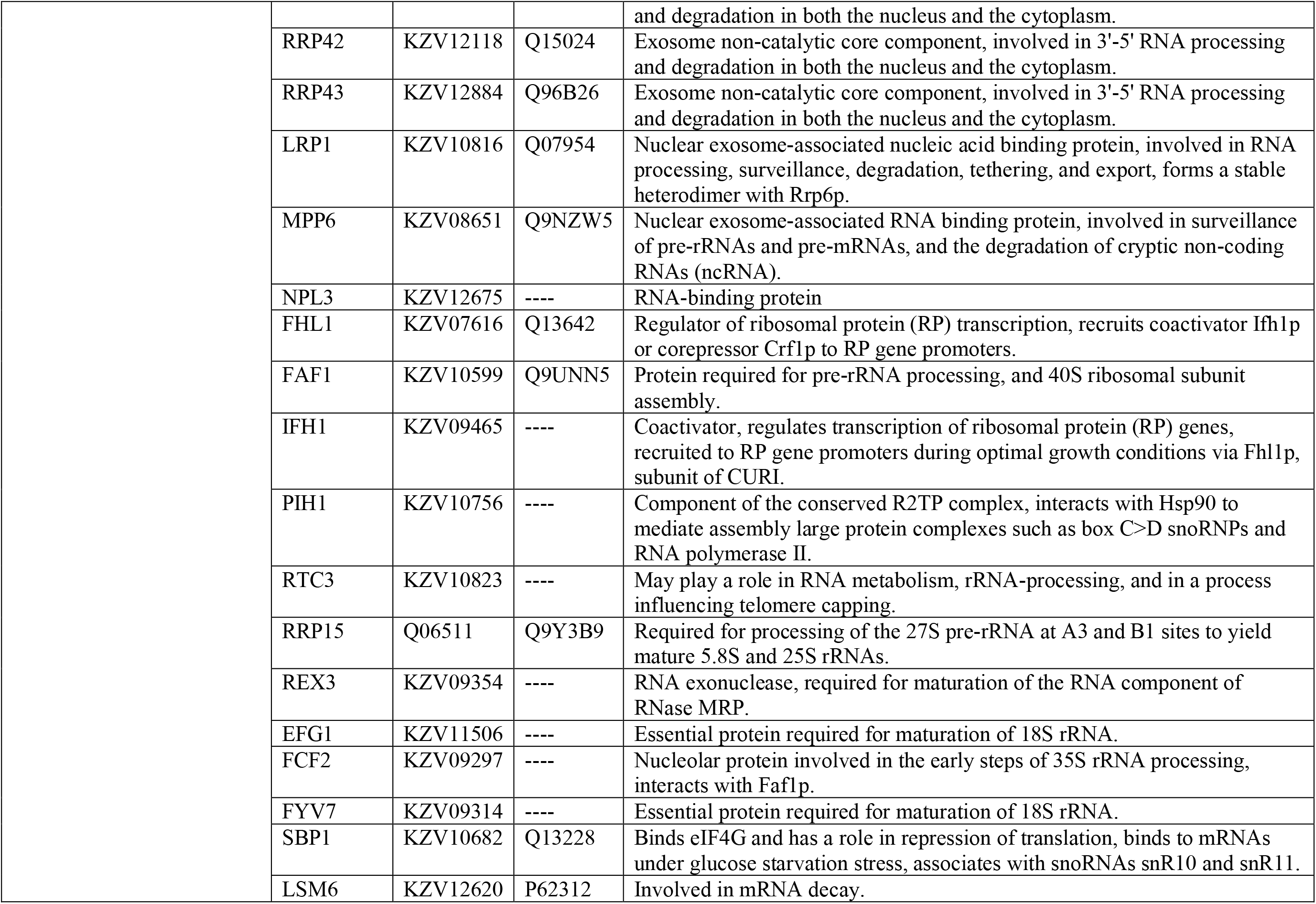
S. cerevisiae ribosome biogenesis proteins with no homologues in E. histolytica.

We further analyzed the repertoire of ribosome biogenesis proteins, to understand the conserved and divergent features of this essential function in *E. histolytica*. We observed that a number of different proteins were represented by a single gene in *E. histolytica* as opposed to multiple genes in *S. cerevisiae* (Table 3). These belonged to protein families with very similar sequences and might have evolved from the same gene by duplication and divergence. There were 42 such protein families and they mostly included helicases, casein kinases, exonucleases and methyltransferases. We also found that the factors required for mitochondrial ribosome biogenesis like MTF1, MTG, GEP3 and MRM1 (217, 218, 219 and 225 in Supplementary Table 1), were not present in *E. histolytica* which could be expected since *E. histolytica* lacks typical mitochondria (54,55). All of the Box C/D snoRNPs and Box H/ACA snoRNPs were present in *E. histolytica* except for a non-core component NAF1(56), and the expression level of snoRNPs was generally high (with a few showing medium expression) (Table 1), suggesting that the process of pre-rRNA modification (methylation and pseudo-uridylation) is well conserved and extensive in *E. histolytica*.

**Table 3:**
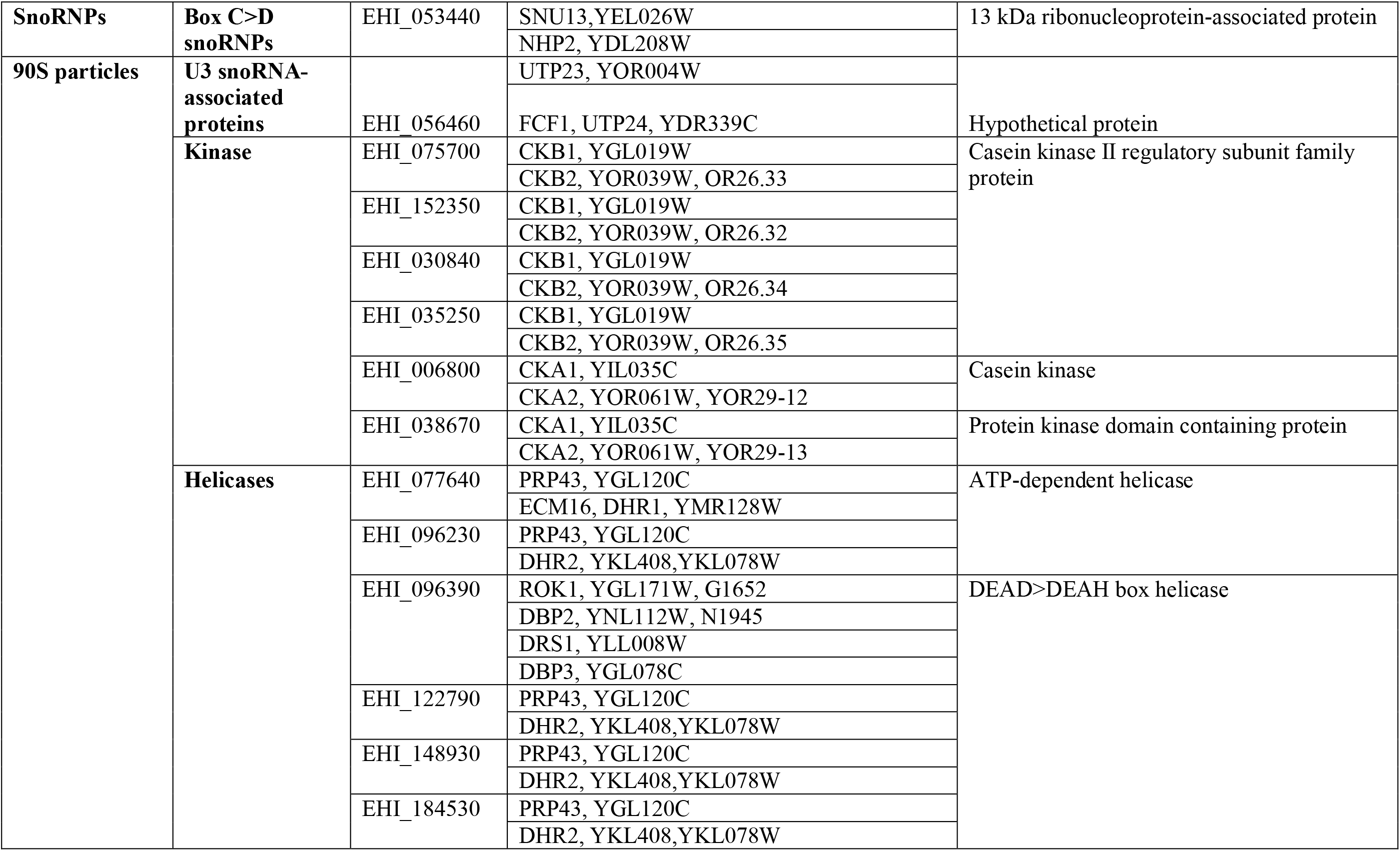
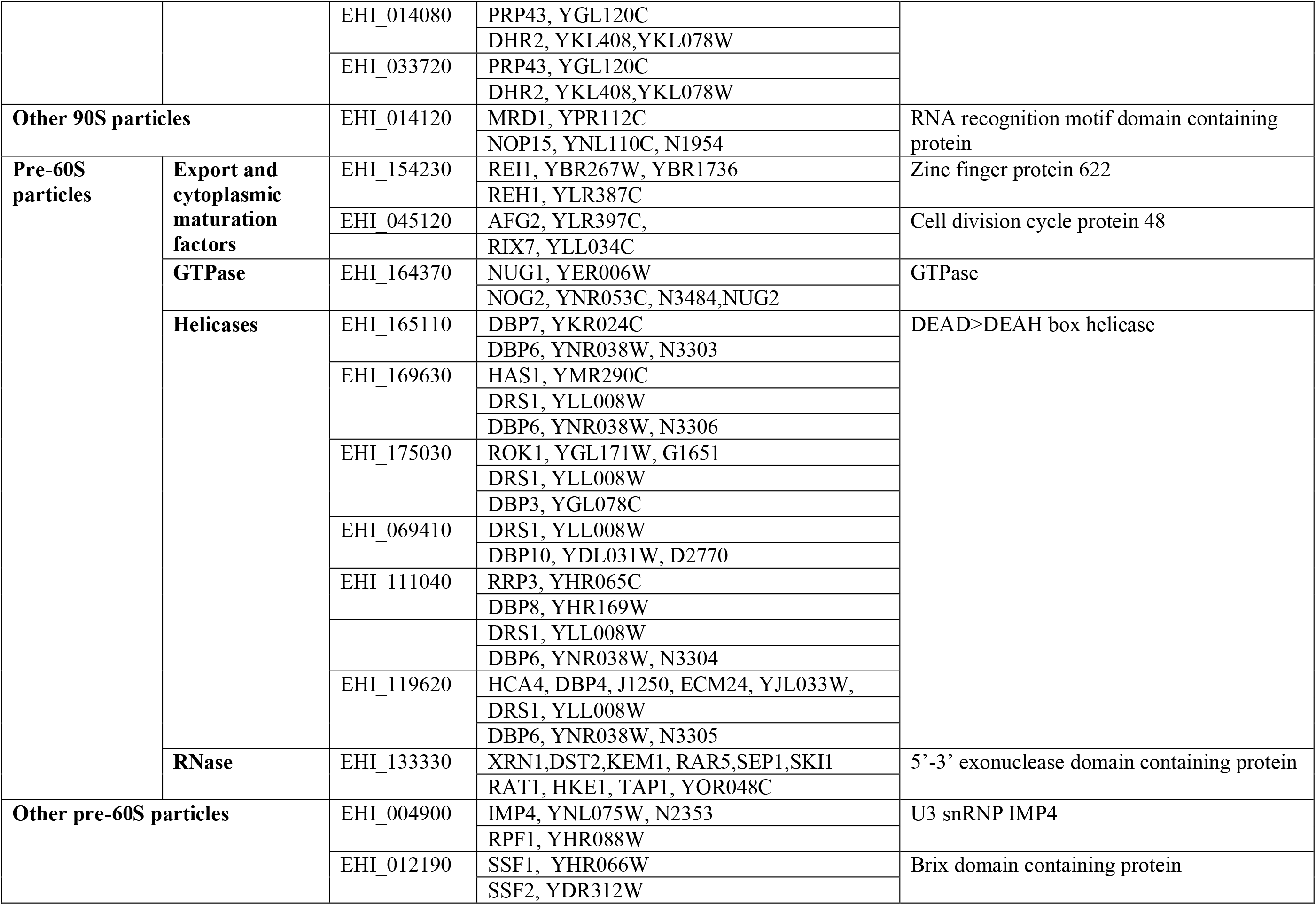
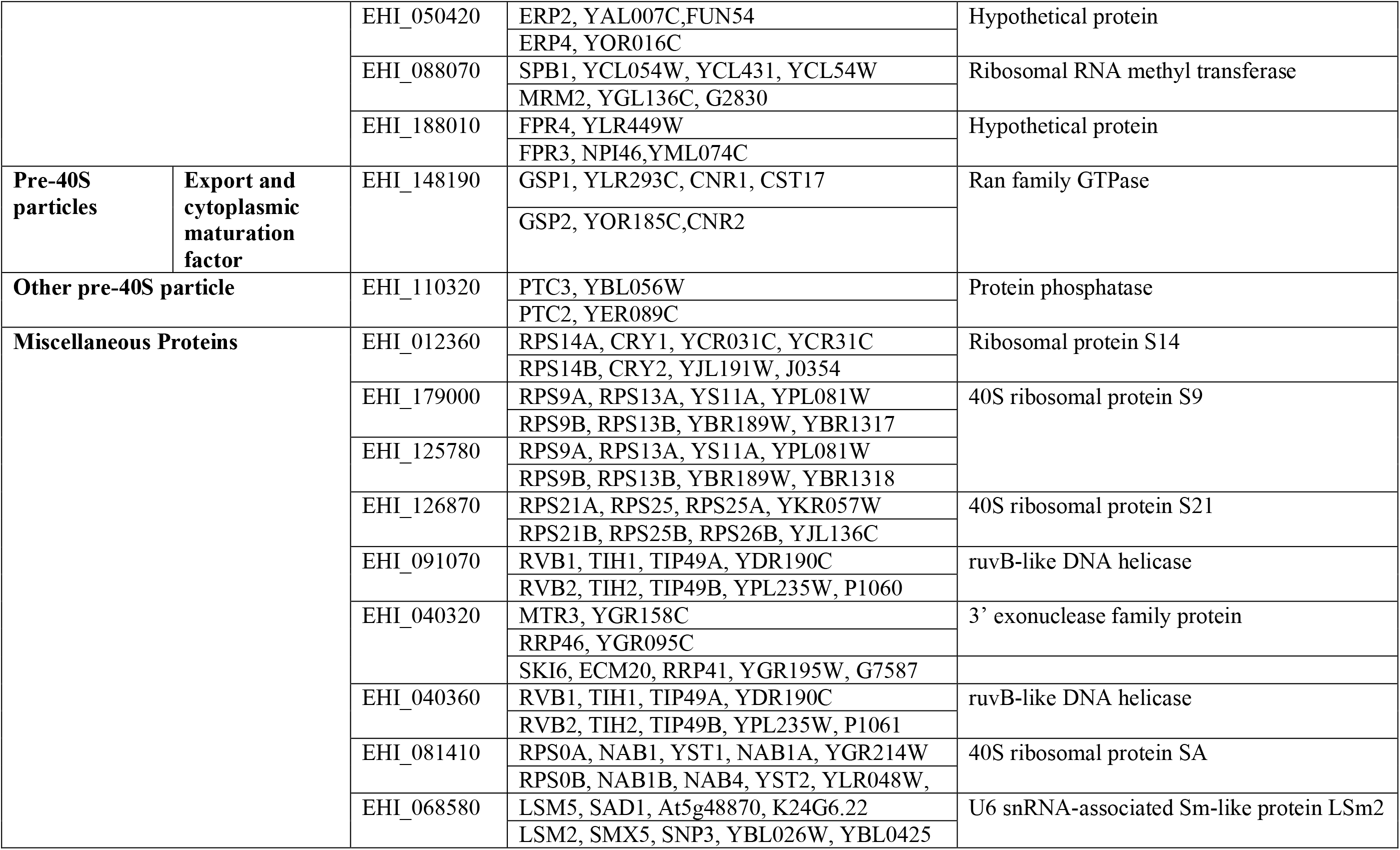
*E. histolytica* genes which match multiple *S. cerevisiae* genes.

Amongst all the factors associated with 90S-, pre-60S- and pre-40S-particles, it was the RNases that were strikingly less conserved in *E. histolytica*, compared to the helicases, GTPases and Kinases. We failed to find any homologue of the RNases RNT1, REX4, NGL2, REX2 and REX1 (Supplementary Table 1.2.4; 1.4; Table 4). The RNases found in *E. histolytica* included a single copy of debranching enzyme DBR1, containing an N-terminal metallophos domain, required for its activity. There were two copies of RAT1 and a single copy of XRN1(57). RAT1 together with its counterpart XRN1 are highly conserved 5’-3’ exoribonucleases. They play a crucial role in gene transcription, RNA processing and RNA surveillance. These are required specifically for regulating mRNA homeostasis by removing processing intermediates, aberrant molecules, and decapped mRNAs, by acting post transcriptionally and co-transcriptionally(58–60). Like *S. cerevisiae*, Humans also possess 8 different RNases (51) most of which were not found in *E. histolytica* (Table 4). It is possible that *E. histolytica* utilizes a limited number of RNases to target a variety of substrates.

**Table 4:**
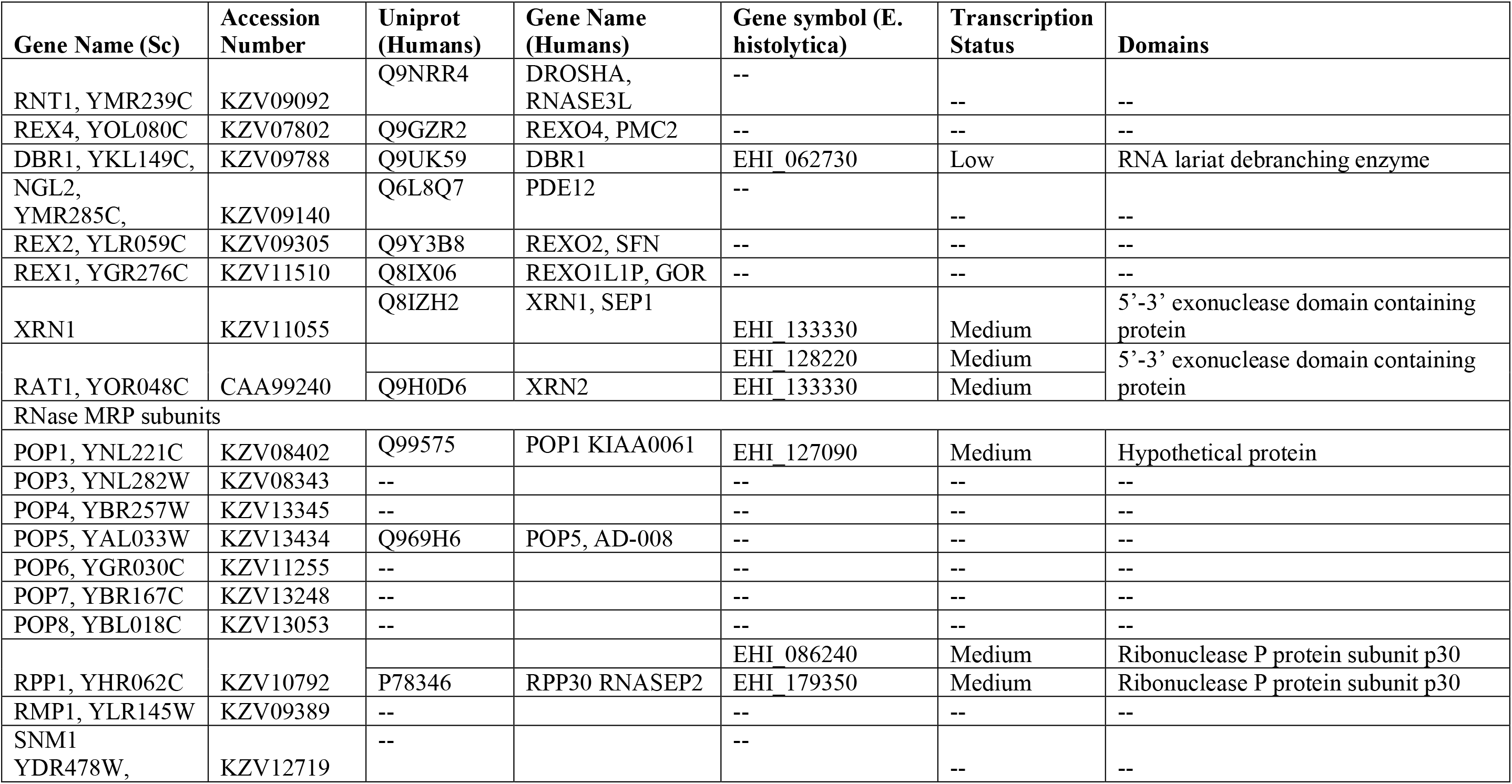
RNases in S. cerevisiae and homologs in human and E.histolytica.

| Gene Name (Sc) | Accession Number | Uniprot (Humans) | Gene Name (Humans) | Gene symbol (E. histolytica) | Transcription Status | Domains |
| --- | --- | --- | --- | --- | --- | --- |
| RNT1, YMR239C | KZV09092 | Q9NRR4 | DROSHA, RNASE3L | -- | -- | -- |
| REX4, YOL080C | KZV07802 | Q9GZR2 | REXO4, PMC2 | -- | -- | -- |
| DBR1, YKL149C, | KZV09788 | Q9UK59 | DBR1 | EHI_062730 | Low | RNA lariat debranching enzyme |
| NGL2, YMR285C, | KZV09140 | Q6L8Q7 | PDE12 | -- | -- | -- |
| REX2, YLR059C | KZV09305 | Q9Y3B8 | REXO2, SFN | -- | -- | -- |
| REX1, YGR276C | KZV11510 | Q8IX06 | REXO1L1P, GOR | -- | -- | -- |
| XRN1 | KZV11055 | Q8IZH2 | XRN1, SEP1 | EHI_133330 | Medium | 5'-3' exonuclease domain containing protein |
| RAT1, YOR048C | CAA99240 | Q9H0D6 | XRN2 | EHI_128220<br>EHI_133330 | Medium<br>Medium | 5'-3' exonuclease domain containing protein |
| RNase MRP subunits |  |  |  |  |  |  |
| POP1, YNL221C | KZV08402 | Q99575 | POP1 KIAA0061 | EHI_127090 | Medium | Hypothetical protein |
| POP3, YNL282W | KZV08343 | -- |  | -- | -- | -- |
| POP4, YBR257W | KZV13345 | -- |  | -- | -- | -- |
| POP5, YAL033W | KZV13434 | Q969H6 | POP5, AD-008 | -- | -- | -- |
| POP6, YGR030C | KZV11255 | -- |  | -- | -- | -- |
| POP7, YBR167C | KZV13248 | -- |  | -- | -- | -- |
| POP8, YBL018C | KZV13053 | -- |  | -- | -- | -- |
| RPP1, YHR062C | KZV10792 | P78346 | RPP30 RNASEP2 | EHI_086240<br>EHI_179350 | Medium<br>Medium | Ribonuclease P protein subunit p30<br>Ribonuclease P protein subunit p30 |
| RMP1, YLR145W | KZV09389 | -- |  | -- | -- | -- |
| SNM1 YDR478W, | KZV12719 | -- |  | -- | -- | -- |

The RNase MRP(RMRP) is another important ribonuclease associated with pre-60S particles (Supplementary Table 1.4). It is a ribonucleoprotein complex of 8-10 protein subunits which are bound to an RNA molecule required for catalytic activity (9). Our data showed very low sequence homology with two RMRP subunits (POP1and RPP1), while the other subunits were absent in *E. histolytica* (Supplementary Table 1.4.3.1). Amongst other protozoan parasites *Giardia lamblia* is reported to have five RMRP protein subunits and *Leishmania* and *Trypanosoma* showed homology with only one subunit (RPP25) (61). The presence of RMRP RNA component is reported in *E. histolytica* (62,63), while it is absent in *Leishmania*, *Trypanosoma* and *Giardia*. RMRP is known to cleave pre-rRNA at the A3 site in the ITS 1, in a manner similar to RNase P (64,65), the RNA component of which acts as a ribozyme (66). Mutations in the RNA component of RMRP are reported to cause cartilage hair hypoplasia (67,68). However, the absence of RNA component in RMRP from a number of parasites, and poor conservation of protein subunits suggests significant evolutionary diversity of this enzyme.

Amongst the proteins associated with 90S particles are the U3 snoRNPs (Supplementary Table 1.2). Previous work has shown that *E. histolytica* has 5 copies of U3snoRNA (69) which associate with conserved U3snoRNPs for modification of pre-rRNA. Of the 30 U3snoRNPs in *S. cerevisiae,* 23 were present in *E. histolytica*. The seven missing components were RRP7, NOP14, UTP3, UTP8, UTP9, UTP16 and UTP30. Of these UTP8, UTP9 and UTP16 were also missing in human. Amongst other proteins in 90S particles we found that 8-10 DEAD/DEAH box helicases and a single GTPase BMS1 was present in *E. histolytica*. Of the twenty-two other 90S factors, eight were missing in *E. histolytica* of which CIC1, CMS1 and NOP6 were also missing in human and were non-essential in *S. cerevisiae*. The other five (KRI1, NOP9, PXR1, RRP36 and TMA23) were present in human while absent in *E. histolytica*.

In the pre-40S particles (Supplementary Table 1.3), of the 20 export and cytoplasmic maturation factors required during the formation of 40S, three factors LTV1, SXM1 and KAP114 were missing in *E. histolytica*, all being present in human. SLX9 and eukaryotic translation initiation factor 3 subunit J (HCR1) were also missing from pre-40S particles. The most important export factor CRM1 known to be responsible for the export of both 60S and 40S subunits to the cytoplasm was present in *E. histolytica* with moderate homology and sequence similarity; however, it is annotated as a hypothetical protein in the database. An important kinase of 60S particle GRC3 (NOL9 in humans), which cooperates with the endoribonuclease LAS1 to process pre-rRNA at site C2 in ITS2 (70), was also missing in *E. histolytica*. However XRN2, which is also reported to play a role in the processing of ITS2, was present (Figure 1a) (71). Protein kinases HRR25 and RIO2 which are known to associate with pre-40S particle were present in *E. histolytica.* HRR25 phosphorylates ribosomal protein RPS3, dephosphorylation of which marks the maturation of 40S particle (72). A number of other accessory factors were missing in *E. histolytica.* Some of the unique features of *E. histolytica*, namely organization of rRNA genes as extrachromosomal circles (41), and location of nucleolus at the nuclear periphery (42), may explain the differences in ribosome biogenesis machinery between *E. histolytica* and *S. cerevisiae*. Our data highlighted that 67% of the 279 *S. cerevisiae* genes involved in ribosome biogenesis could be found in *E. histolytica* with a high degree of sequence conservation, implying the use of most of the conserved pathways found in *S. cerevisiae*. It will be interesting to see the impact of the missing 33% genes (some of which may have been missed due to poor homology) on alternative strategies of ribosome biogenesis employed by *E. histolytica*, and possibly by other protists.

**Figure 1:**
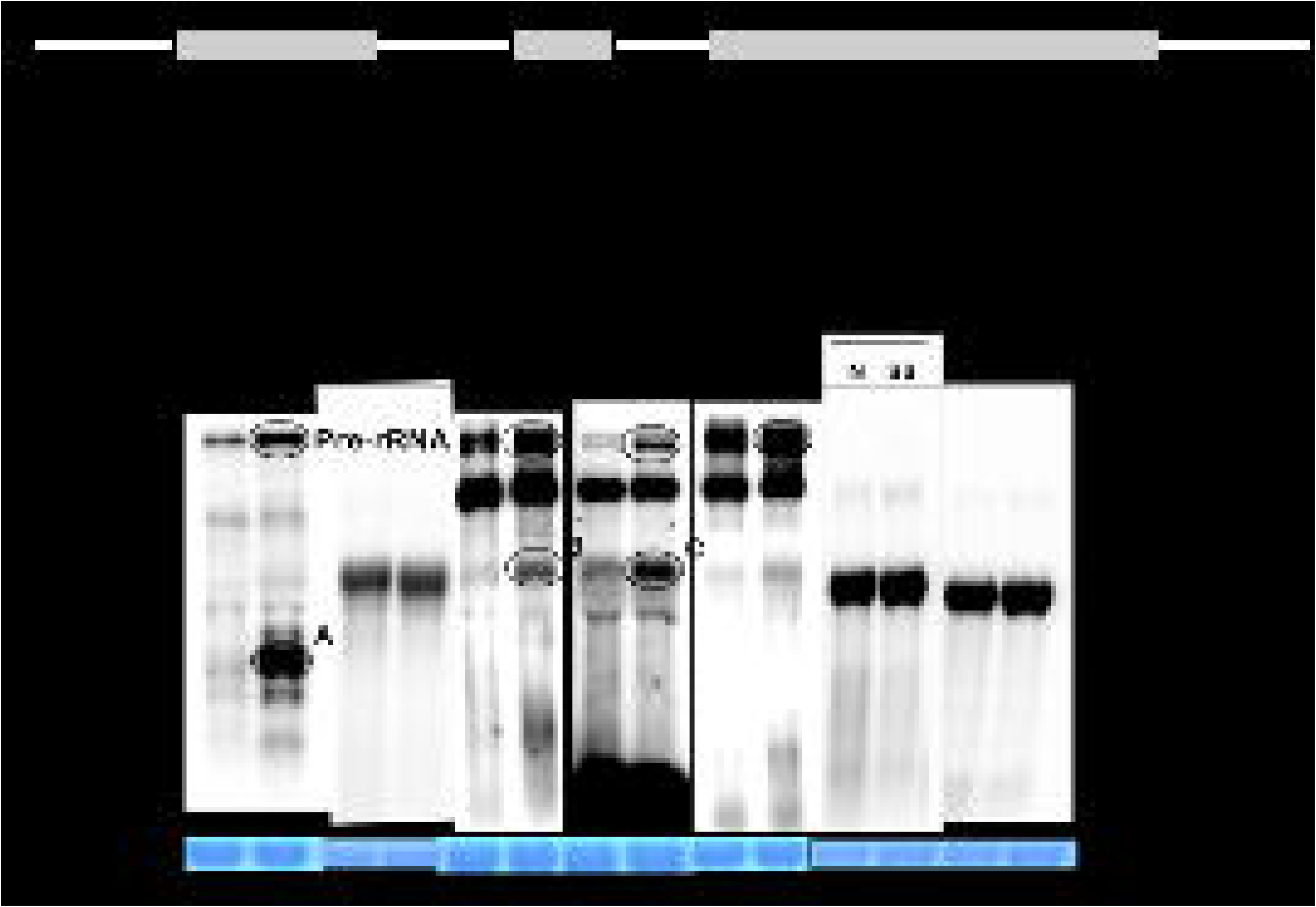
(a) Schematic linear view of rDNA transcription unit indicating Sc processing sites. The dotted arrows indicate the sites present in *E. histolytica.* The size of the 5’-ETS (2.672 kb), 18S (1.921 kb), ITS1 (149 bp), 5.8S (150 bp), ITS2 (123 bp), 28S (3.544 kb) and IGS (5.183 kb) accounts for the 14 kb rDNA of *E. histolytica*. (b) Northern hybridization of total RNA from *E. histolytica* cells grown under normal (N) and serum starved (SS) conditions. The probes used are indicated. The lanes with 18S and 28S probes were exposed very briefly and show the position of the mature rRNAs. 28S rRNA in *E. histolytica* has an internal nick and two probe sets were used as indicated. The bands accumulated in SS cells are marked in circle.

### Effect of growth stress on transcription of pre-rRNA processing and ribosome biogenesis-genes in *E. histolytica*

Stress signals cease the transcription of rRNA genes in most model organisms. The demand for ribosomes is reduced in response to various environmental stresses e.g. serum starvation (73), inhibition of protein synthesis by cycloheximide treatment, and oxidative stress (74–79). This growth-dependent regulation of rRNA synthesis is conserved from bacteria to vertebrates (73). Growth stress induced by serum starvation in *E. histolytica* leads to the accumulation of unprocessed pre-rRNA, and 5’-ETS intermediates, which are otherwise rapidly degraded during normal growth conditions (46). We have earlier shown that the exosomal subunit RRP6 is lost from the nucleus during serum starvation, which could lead to the accumulation of 5’-ETS sub fragments (47). To obtain a more comprehensive picture of the defects in pre-rRNA processing during growth stress in *E. histolytica* we looked for the accumulation of other intermediates during serum starvation, using probes from each segment of pre-rRNA, in a northern hybridization analysis. We observed an accumulation of different intermediates in *E. histolytica* with all the probes. The 5’-ETS probe, showed accumulation of 0.7 -0.9 kb fragment while the ITS1, 5.8S and ITS2 probes showed accumulation of a ∼2 kb intermediate (Figure 1b). These data imply a more generalized defect in pre-rRNA processing during serum starvation. This could be due to the cumulative effect of loss of activity of a number of proteins, including endo- or exonucleases, required for the cleavage and trimming of pre-rRNA, and RNA helicases, the role of which in accumulation of pre-rRNA intermediates is well known (80).

A number of check points prevent entry of immature ribosomes into the translating pool (8,11). The discovery of ribosomopathies, due to mutations in genes encoding ribosome biogenesis factors, has shed new light on the link between ribosome biogenesis and cell fate (81). Defects in pre-rRNA processing, nucleolar organization and ribosomal subunit accumulation have been detected in the cells from patients suffering from ribosomopathies. To explore the possible factors responsible for the defects in pre-rRNA processing during growth stress in *E. histolytica*, we analyzed the transcriptome of trophozoites grown in normal conditions and after 24hrs of serum starvation. Differential RNA expression analysis of all the ribosome biogenesis proteins present in *E. histolytica* led us to identify a number of possible factors that could interfere in pre-rRNA processing. The expression values of all the listed genes in normal and serum starved growth conditions are represented as heat map in Figure 2a. Of the 253 genes involved in pre-rRNA processing in *E. histolytica,* 40 genes showed differential expression (>1.5-fold change) of which 22 were upregulated and 18 were downregulated (Table 5, 6 respectively). The upregulated genes included three HSPs, four helicases, casein kinase, mRNA decay protein, and cytoplasmic export protein, NMD3. Casein Kinase showed the highest fold change of 3.56. NMD3 upregulation indicates the initiation of mRNA decay pathway during growth stress. RNA helicases function at many steps, including snoRNA unwinding after pre-rRNA modification (82) (Table 5, 6). The downregulated transcripts included H/ACA snoRNP complex, NEP1 methylase, NMD4 and the exosome component 10 (RRP6) which has a 3’-5’ exonuclease activity. It is involved in 5’-ETS degradation and 5.8S rRNA processing, and our earlier work has shown that it is down regulated and is lost from nuclei during serum starvation (47,69). We also demonstrated that EhRRP6 acts as a stress sensor, as upregulating EhRRP6 protected the cells against stress while its downregulation hampered cell growth. Casein kinase in yeast is reported to regulate a switch between productive and non-productive pre-rRNA processing pathways post-transcriptionally during stress (83). Downregulation of NEP1 methylase has been shown to block pre-rRNA processing at sites A0, A1 and A2 (84). Thus, incomplete pre-rRNA modification, non-productive pre-rRNA processing, and the inefficient process of RNA intermediate degradation, along with other above-mentioned factors, could together or individually lead to the accumulation of unprocessed intermediates.

**Figure 2:**
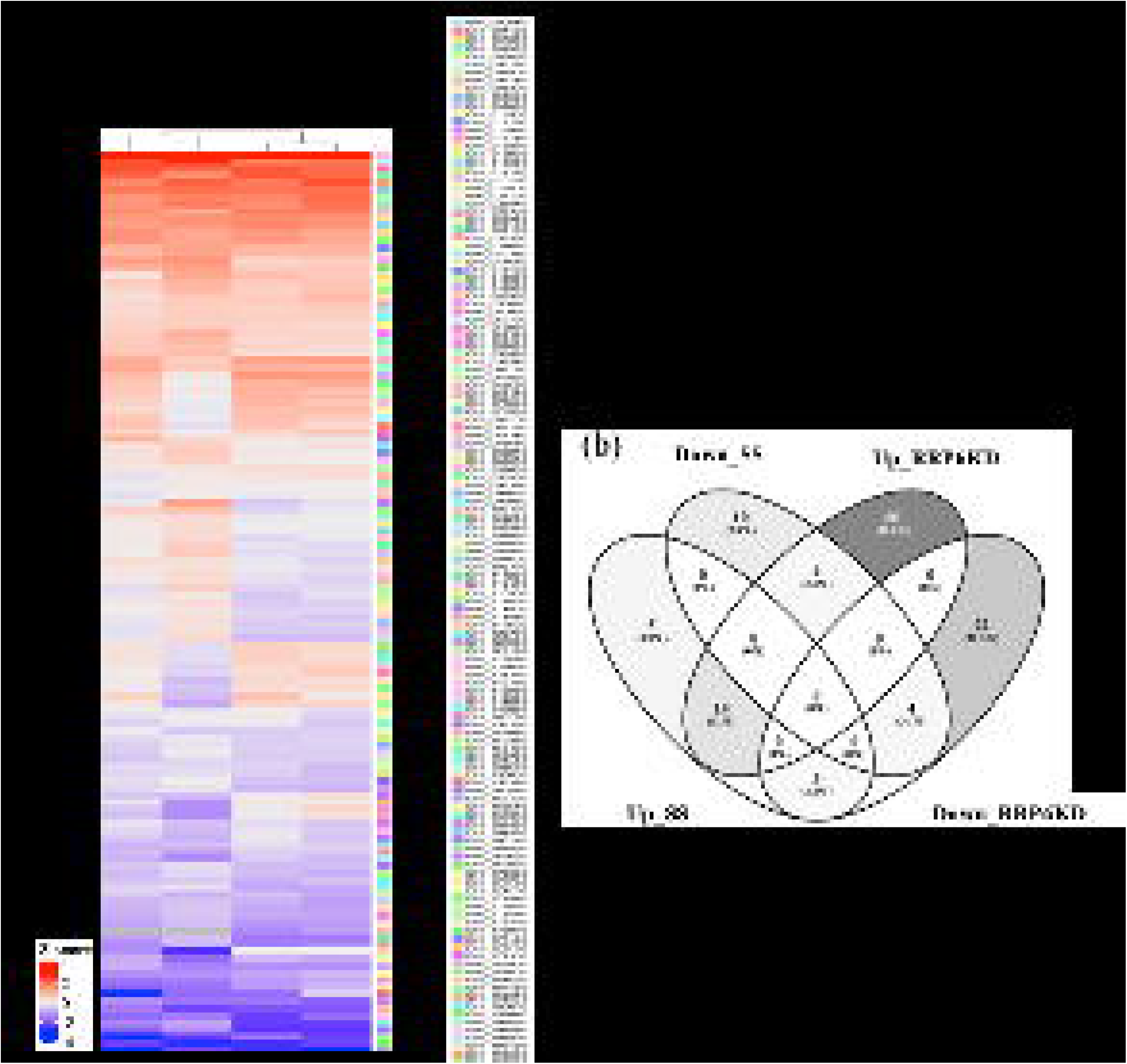
(a) Heat map showing the log_2_ expression of all ribosome biogenesis factors in *E. histolytica* during TOC, Rrp6KD, normal and serum starved growth conditions. Red color depicts high expressing genes and blue color depicts low expressing genes. (b) Venn diagram shows an overlap of the differentially expressed genes in all the four conditions. Down_SS-Downregulated in SS; Up_SS-Upregulated in SS; Down_RRP6KD-Downregulated in RRP6KD; Up_RRP6KD-Upregulated in RRP6KD.

**Table 5:**
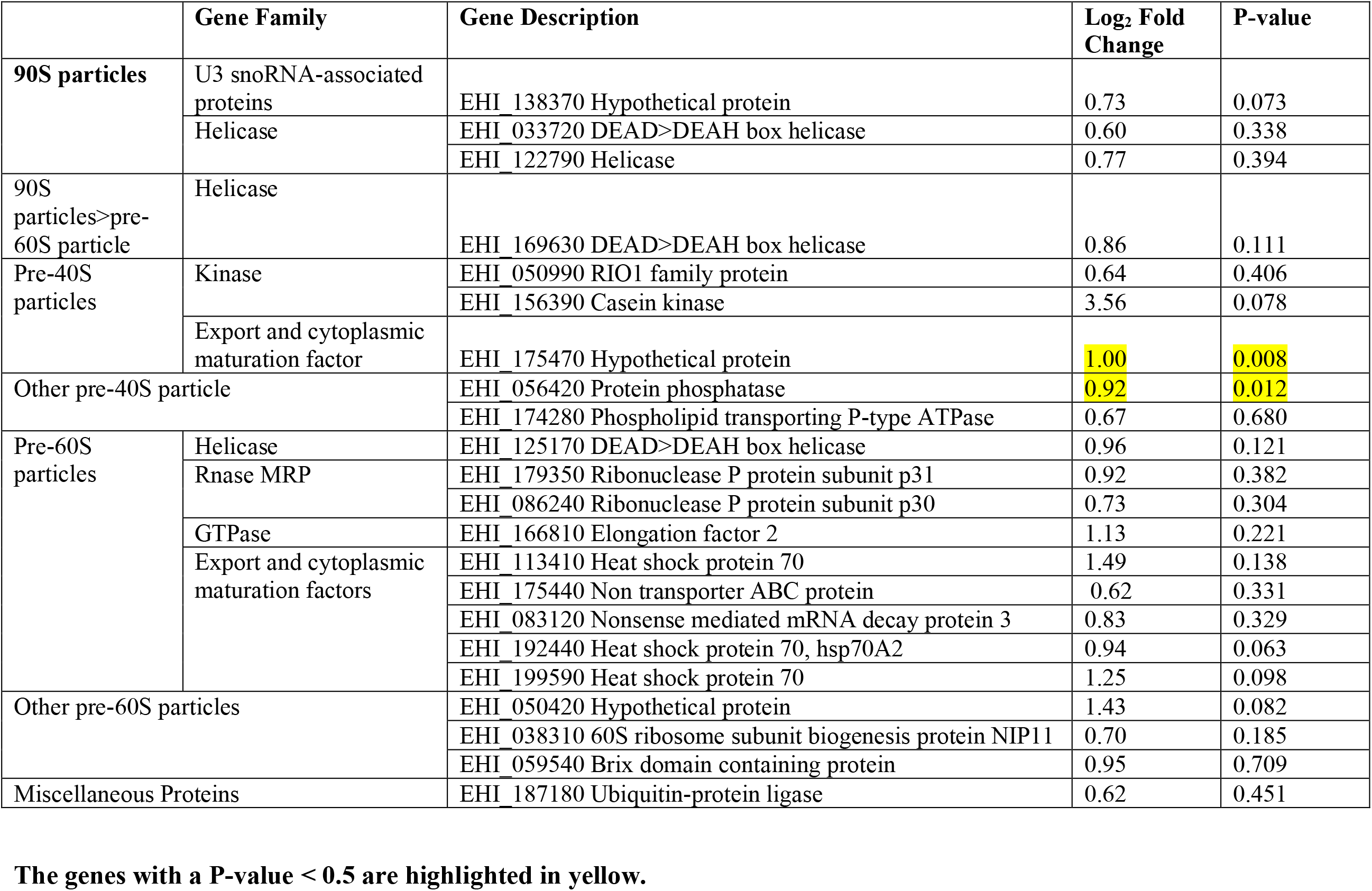
Up regulated (>1.5-fold change) genes during serum starvation as compared to normal.

**Table 6:** Down regulated (>1.5-fold change) genes during serum starvation as compared to normal.

|  | Gene family | Gene Description | Log<sub>2</sub> Fold Change | P-value |
| --- | --- | --- | --- | --- |
| SnoRNPs | Box H>ACA snoRNPs | EHI_001850 H>ACA ribonucleoprotein complex subunit 2-like protein | -0.60 | 0.022 |
|  |  | EHI_102280 H>ACA ribonucleoprotein complex subunit 2-like protein | -0.60 | 0.022 |
| 90S particles | U3 snoRNA associated proteins | EHI_179100 Hypothetical protein | -1.13 | 0.195 |
|  |  | EHI_012960 Hypothetical protein | -0.63 | 0.574 |
|  |  | EHI_198890 Hypothetical protein | -0.61 | 0.469 |
| Other 90S particles |  | EHI_033750 Hypothetical protein | -0.61 | 0.168 |
|  |  | EHI_044610 Ribosome biogenesis protein NEP1 | -1.56 | 0.025 |
| Pre-40S particles | Export and cytoplasmic maturation factor | EHI_164410 Hypothetical protein | -0.60 | 0.524 |
|  |  | EHI_001120 Hypothetical protein | -0.80 | 0.066 |
|  |  | EHI_100490 Hypothetical protein | -0.63 | 0.050 |
| Pre-60S particles | Helicase | EHI_134610 DEAD>DEAH box helicase | -0.79 | 0.209 |
|  | GTPase | EHI_164370 GTPase | -1.03 | 0.111 |
|  | Export and cytoplasmic maturation factors | EHI_052860 Heat shock protein 70 | -0.62 | 0.136 |
|  |  | EHI_179200 Hypothetical protein | -0.65 | 0.475 |
|  |  | EHI_104460 Nonsense mediated mRNA decay protein 4 | -1.23 | 0.476 |
| Other pre-60S particles |  | EHI_105790 Nucleolar complex protein 2 homolog | -0.65 | 0.572 |
| Miscellaneous Proteins |  | EHI_021400 Exosome component 10 | -0.77 | 0.444 |
|  |  | EHI_104570 Ubiquitin ligase | -1.75 | 0.171 |

### Effect of EhRRP6 downregulation on expression of ribosome biogenesis factors in *E. histolytica*

The downregulation of exonuclease RRP6 and its loss from the nucleus during serum starvation (47) led us to investigate the global effect of RRP6 silencing on *E. histolytica* transcriptome compared to the changes observed in serum starvation. The heat map in Figure 2a represents the expression values of all the ribosomal genes in TOC, RRP6 downregulation, normal and SS growth conditions. Of the ribosome biogenesis factors present in *E. histolytica,*102 genes showed >1.5fold change in EhRRP6 silenced cells as compared to 40 genes in serum starved conditions.

78 genes were upregulated in RRP6-silenced cells, of which 63 were significant (p-value < 0.05), 14 of these 78 were also upregulated in serum starved cells, of which only two were significant in both datasets (Figure 2b, Table 7). These two proteins were protein phosphatase (PTC2) and non-transported Abc protein ARB1. Along with the snoRNPs the other upregulated factors included helicases, GTPase, elongation factors, HSPs, cell cycle proteins, other components of exosome complex and RNase MRP. Interestingly ARB1 associates with ribosomes and has been proposed to serve as a mechanochemical ATPase, stimulating multiple steps in 40S and 60S ribosome biogenesis. Its depletion from *S. cerevisiae* led to slower cleavage at A0, A1 and A2 sites in pre-rRNA (85). Its transcriptional up regulation under conditions where pre-rRNA processing has slowed down could indicate its alternate roles in *E. histolytica*, or that the gene may be post-transcriptionally regulated, as is the case with RPs in *E. histolytica* (86).

**Table 7:**
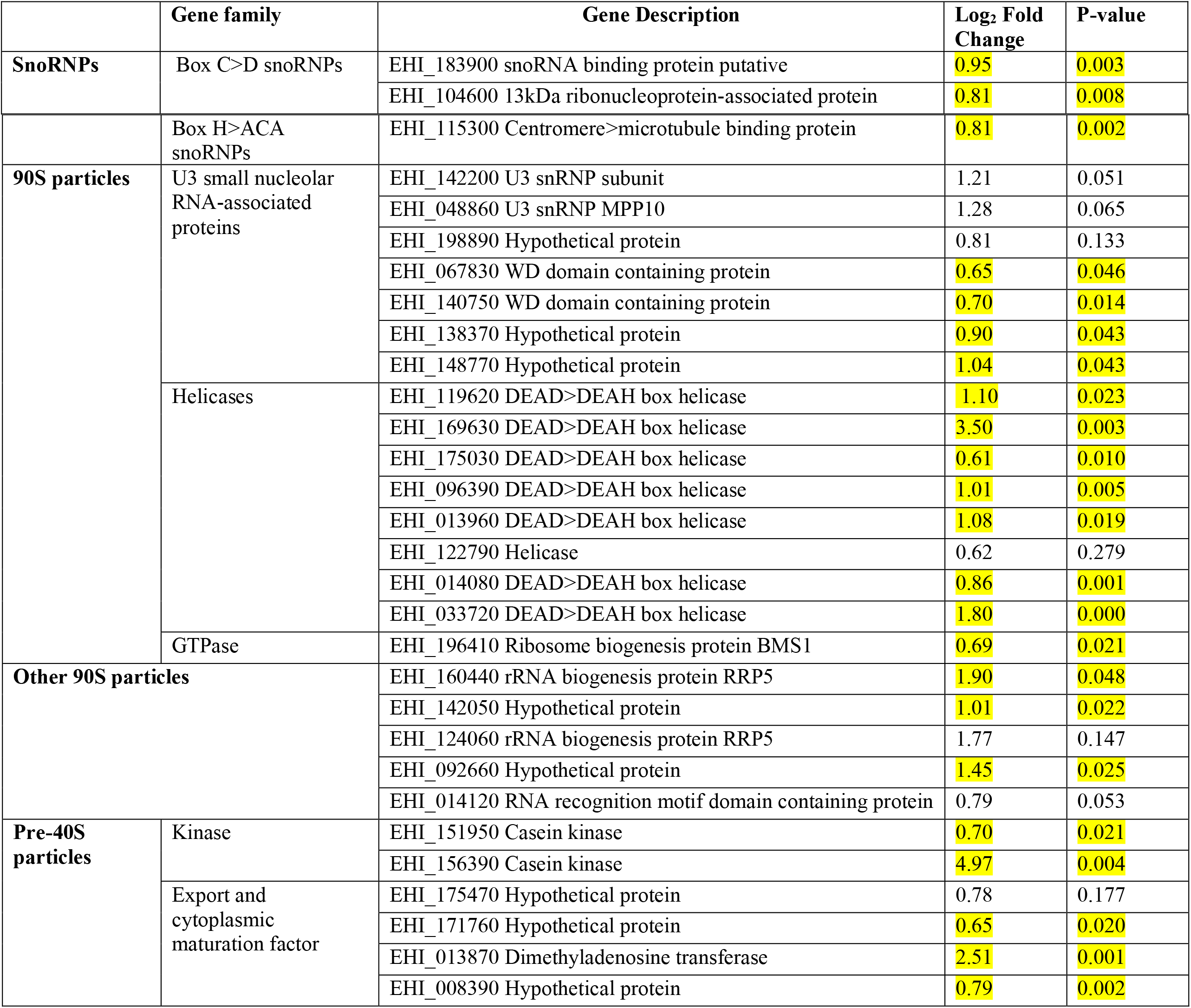
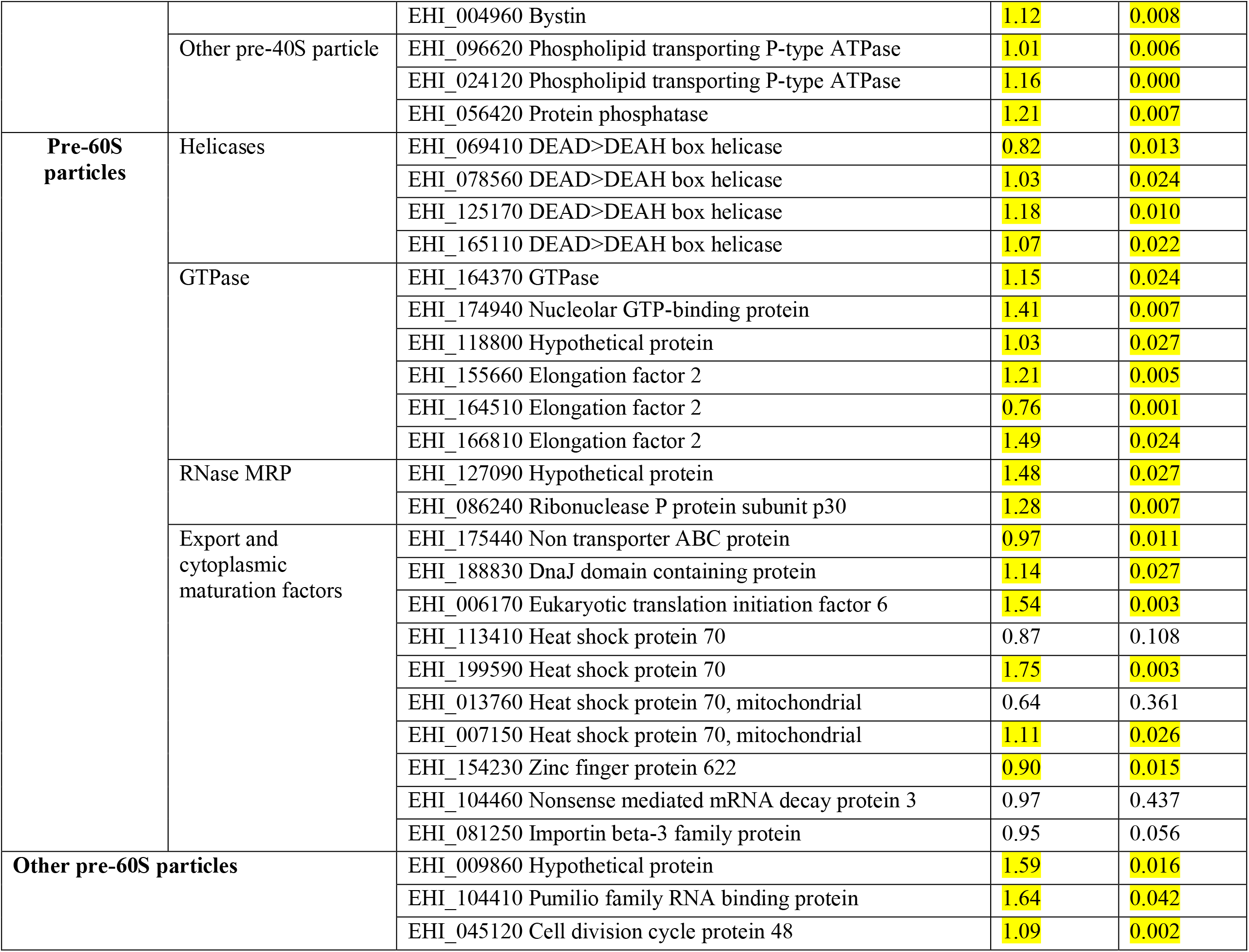
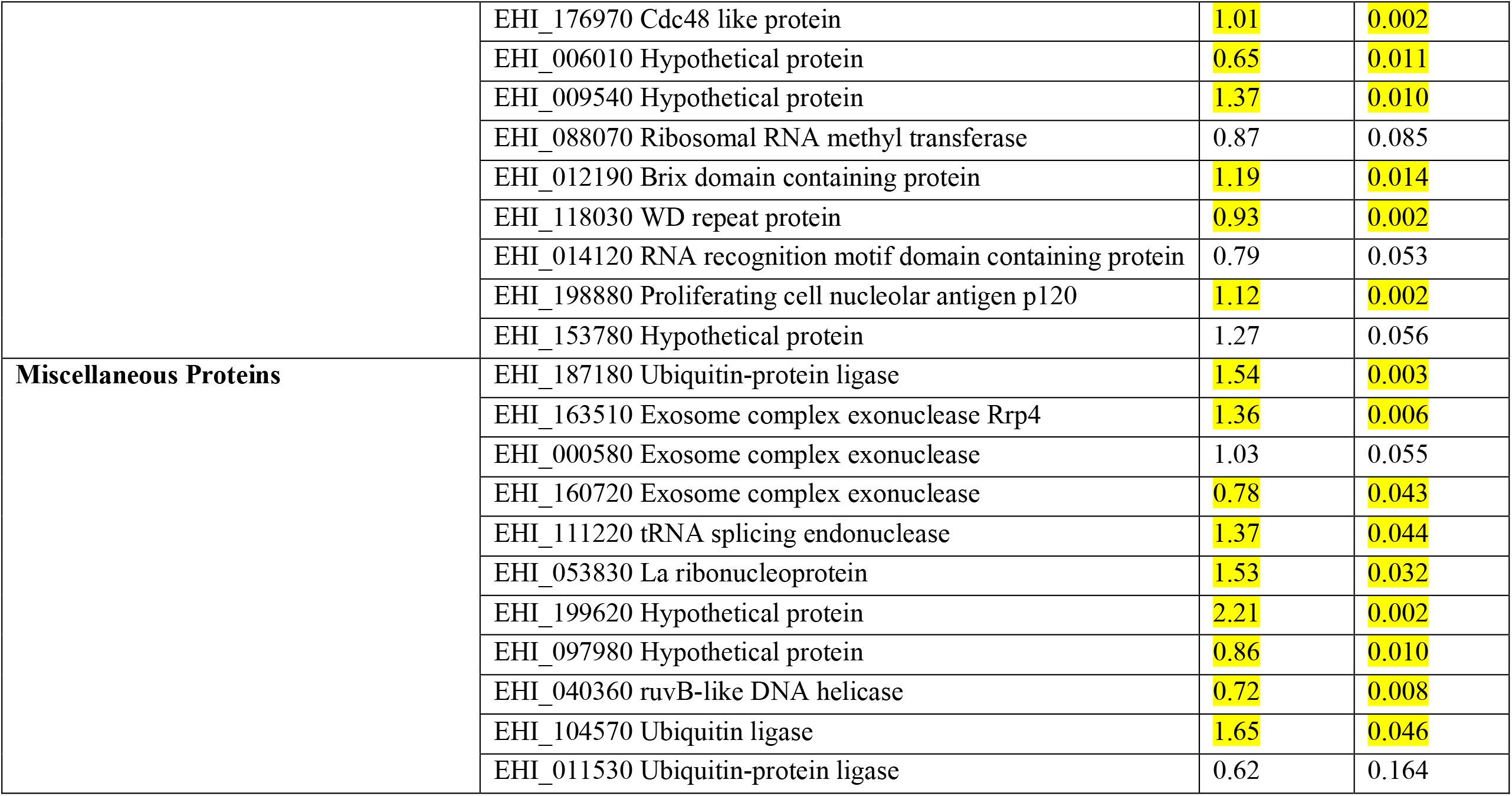
Up-regulated genes (>1.5-fold change) in Rrp6KO as compared to TOC.

34 genes were downregulated in RRP6-silenced cells, with 14 genes having significant p-values (Table 8). These included helicases, HSPs, casein kinases, Box H/ACA snoRNPs and U3 snoRNP subunits, TSR4 (a programmed cell death protein) and LSM-domain containing proteins. Four of these genes were downregulated in serum starved cells as well, with two having significant p-values. These genes, downregulated in both datasets, were NEP1 and TSR3. The latter is annotated as a hypothetical protein in *E. histolytica*. Interestingly, both of these genes are involved in chemical modification of the hypermodified nucleotide N1-methyl-N3-aminocarboxypropyl(acp) pseudouridine located on 18S rRNA next to the P-site tRNA (87). This highly conserved base modification in eukaryotes is completed in three steps. In the first step uridine is converted to pseudouridine guided by a snoRNP. In the second step NEP1 methyltransferase adds a methyl group donated by S-adenosylmethionine (SAM). In the third step, which takes place in the cytoplasm, TSR3 adds the APC group from SAM to this nucleotide. This is a crucial step in 18S rRNA maturation. TSR3 downregulation results in accumulation of 20S rRNA precursor, along with accumulation of full-length pre-rRNA (35S) in yeast and (47S) in human. NEP1 also has a crucial role, and its downregulation results in blockage of pre-rRNA processing at sites A0, A1 and A2 (84). Our data suggest that these proteins are crucial in *E. histolytica* as well, and their expression could be regulated in a common pathway that is responsive both to serum stress and RRP6 downregulation.

**Table 8:**
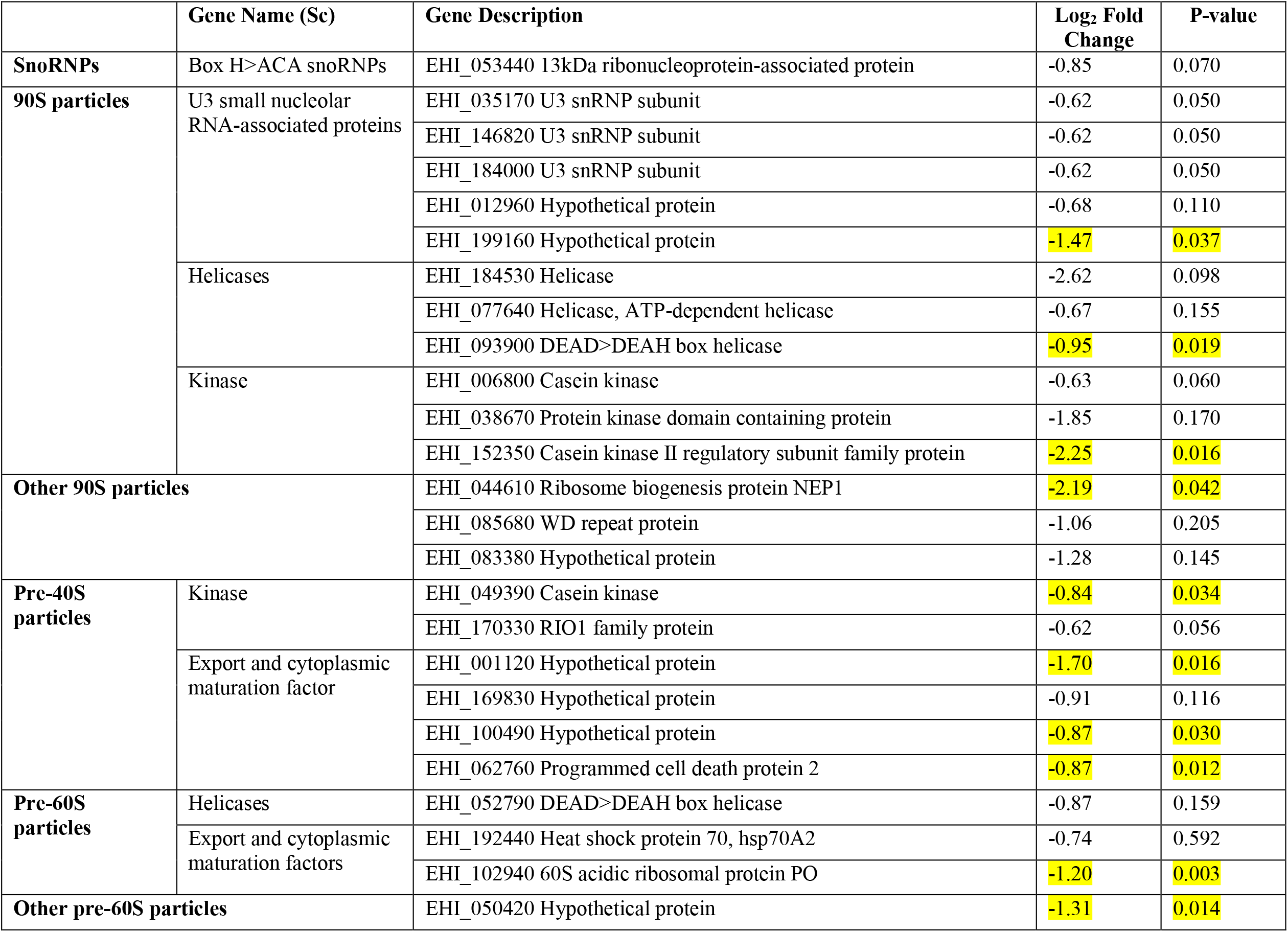
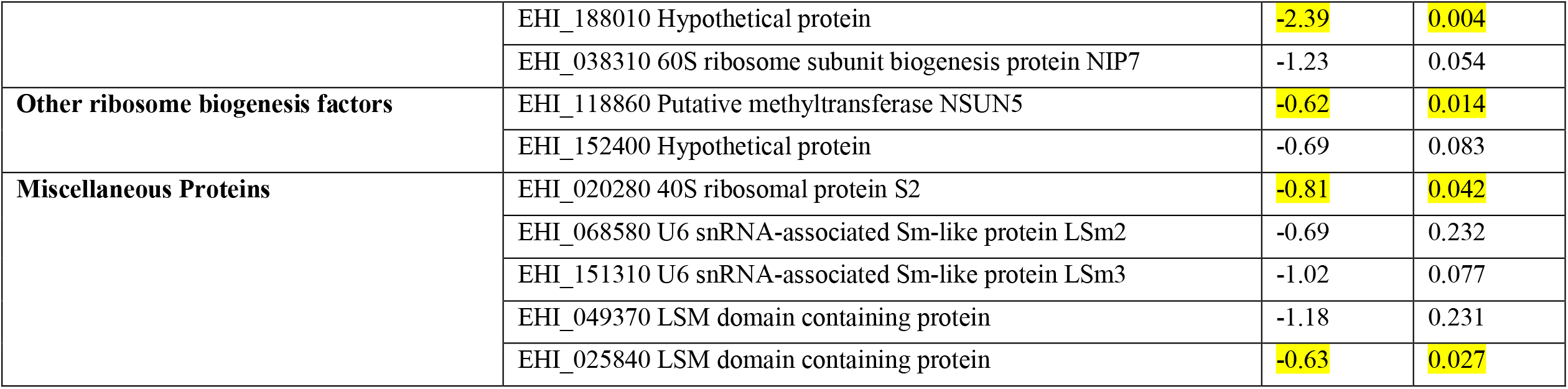
Down regulated genes (>1.5-fold change) in Rrp6KO as compared to TOC.

In conclusion, our data reveal the multiple factors that simultaneously lead to the inhibition of pre-rRNA processing and accumulation of intermediates during growth stress (Figure 3), thus effectively inhibiting ribosome biogenesis by the non-availability of mature rRNAs. Of special interest are the genes affected in a similar manner by both serum starvation and RRP6 down regulation. This work opens up further avenues for detailed molecular investigation of ribosome biogenesis in *E. histolytica*.

**Figure 3:**
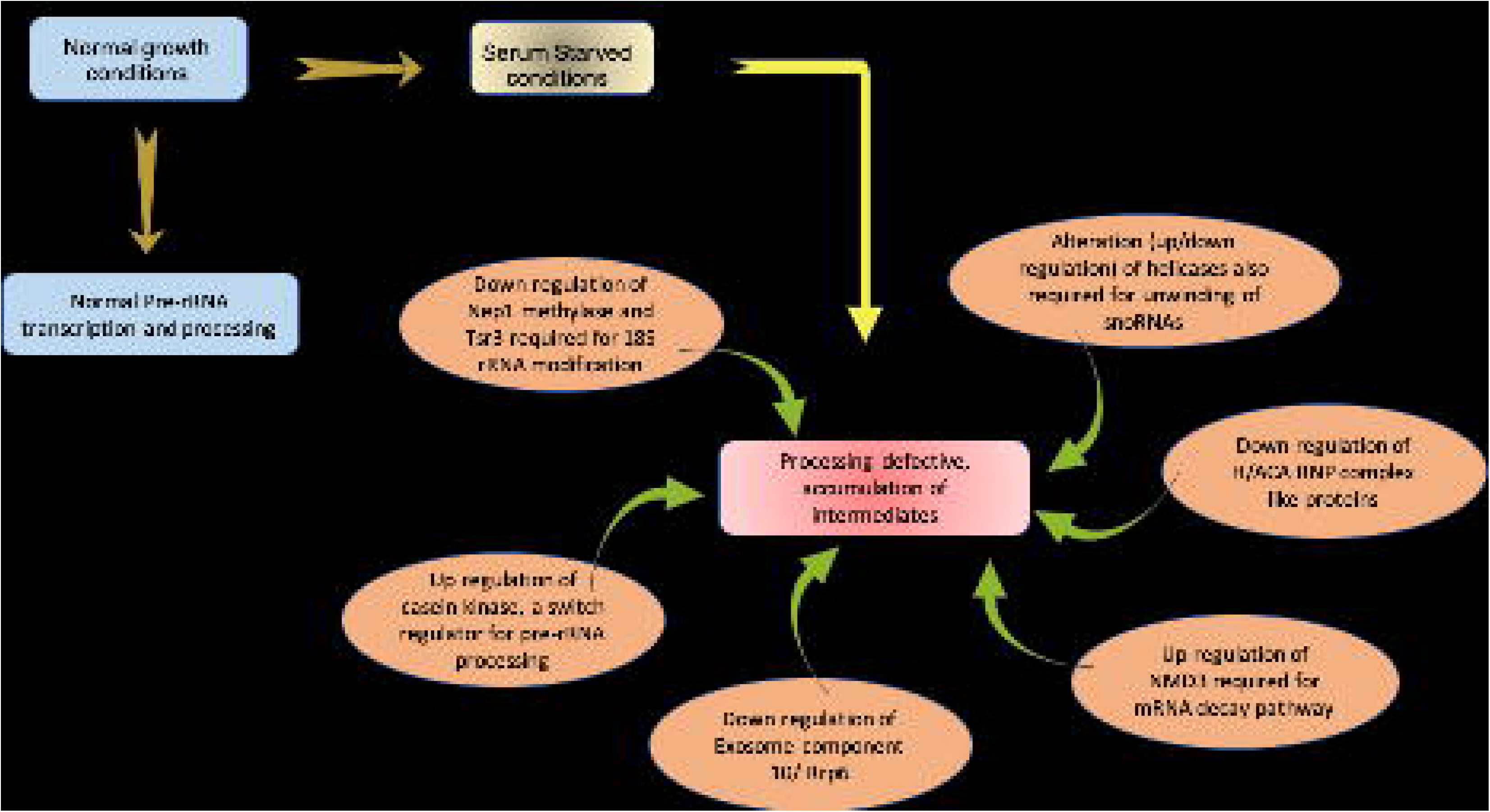
Model depicting factors regulating pre-rRNA processing and accumulation of intermediates during growth stress

## Methods

### Computational identification of *E. histolytica* ribosome biogenesis and pre-rRNA processing proteins

*S. cerevisiae* was used as a reference organism and homologous proteins involved in ribosome biogenesis and pre-rRNA processing in *E. histolytica* were searched. A list of proteins already known to be the part of ribosome biogenesis was compiled using *Saccharomyces* Genome Database (SGD) by applying Gene Ontology Term: ‘rRNA processing’ and ‘Ribosome biogenesis’. The enlisted proteins were further filtered using ‘UniProt’ as Source. Proteins were cross checked in KEGG (Kyoto Encyclopaedia of Genes and Genomes) PATHWAY Database using Keyword: ‘Ribosome biogenesis in eukaryotes’ and filtered keeping ‘*S. cerevisiae*’ as the concerned organism. Proteins were annotated using ‘Function’ from UniProt or ‘Description’ from SGD. Homologous proteins in *E. histolytica* were obtained using PANTHER (Protein Analysis THrough Evolutionary Relationships), AmoebaDB and EggNOG (evolutionary genealogy of genes: Non-supervised Orthologous Groups) databases. BLASTP was performed against respective protein sequence of *E. histolytica* obtained from AmoebaDB restricting the search only to “*Entamoeba histolytica* HM-1:IMSS (taxid:294381)” to obtain an E-value, score, coverage, maximum identity and domains present. The proteins were functionally classified using the KEGG classification. The accession number and annotation were obtained from NCBI, protein database. The expression level of respective genes was extracted from the transcriptome data.

### Cell culture and growth conditions

Trophozoites of *E. histolytica* strain HM-1:IMSS were axenically maintained in TYI-S-33 medium supplemented with 15% adult bovine serum (Biological industries, Israel), Diamond’s Vitamin mix, Tween 80 solution (Sigma–Aldrich) and antibiotics (0.3 units/ml penicillin and 0.25 mg/ml streptomycin) at 35.5^0^C (88). For serum starvation, medium from early to mid-log phase grown trophozoites (48 hrs) was replaced with TYI-S-33 medium containing 0.5% adult bovine serum and incubation continued for 24hrs.

### RNA isolation and Transcriptome analysis

Cells were harvested from normal, serum starvation, TOC and Rrp6-depleted conditions (47). Total RNA from ∼5 × 10^6^ cells were purified using TRIzol reagent (Invitrogen) according to the manufacturer’s instructions. Total RNA, from two biological replicates of each sample was used for selection of polyA plus RNA and library preparation was done after oligo (dT) selection. RNA-Seq libraries were generated and subjected to paired-end sequencing on the Illumina HiSeq2500 (v3 Chemistry) platform (52). The pre-processed reads were aligned to the *E. histolytica* (HM1:IMSS) genome for which the gene model was downloaded from AmoebaDB (http://amoebadb.org/common/downloads/release-27/EhistolyticaHM1IMSS/gff/data/). The alignment was performed using Tophat program (version 2.0.11) with default parameters. The RSEM program (version 1.3.0) was used for estimating expression of the genes and transcripts(89). The differential gene expression analysis was performed using cuffdiff program of cufflinks package with default settings to analyze the difference between normal and serum starved cells. Log_2_ fold change was set as a cut-off for differential expression. Real time qRT PCR was done to validate the data (52). Heatmap was prepared using ggplot2 in R Studio. The Venn diagram was plotted using Venny^2.1^ (https://bioinfogp.cnb.csic.es/tools/venny/).

### Northern blotting

For Northern blot analysis 10 μg of total RNA was resolved on a 1.2% formaldehyde-agarose gel in gel running buffer (0.1 M MOPS (pH 7.0), 40 mM sodium acetate, 5 mM EDTA (pH 8.0)) and 37% formaldehyde at 4 V/cm. The RNA was transferred on to GeneScreen plus R membrane (PerkinElmer). [α-^32^P]dATP-labeled probe was prepared by random priming method using the DecaLabel DNA labeling kit (Thermo Scientific). Hybridization and washing conditions for RNA blots were as per the manufacturer’s instructions.

## Competing interests

The authors declare no competing interests.

## Funding

This work was supported by DAE-BRNS and INSA fellowship to SB, Council of Scientific and Industrial Research fellowship to SN and SSS.

## Authors’ contributions

SN and SB were involved in the study design. SN, SSS, DK, YP, AM conducted the study. SN and SB analyzed and interpreted the data. SN and SB drafted the manuscript. All authors read and approved the final manuscript.

## Acknowledgement

We thank Sneha Bhat, Central University of Punjab, Bathinda, for her contribution in RNA-Seq analysis.

## Notes

### Competing Interest Statement

The authors have declared no competing interest.

